# UcTCRp: a TCRβ-based framework for quantitative MAIT- and iNKT-associated repertoire-state profiling

**DOI:** 10.64898/2026.05.25.727598

**Authors:** LinLin Chen, Yueyi Li, Shiwen Shan, Kaixuan Wang, Chao Feng, Yunsheng Dou, Qihang Xu, Linru Cai, Hao Wang, Haiyang Wang, Xiaochen Bo, Jian Zhang

## Abstract

MAIT and iNKT cells are conventionally identified using invariant or semi-invariant TCRα chains, antigen-loaded tetramers, or transcriptomic phenotypes. These requirements limit their detection in public and clinical immune-repertoire datasets that contain only TCRβ sequences. Here we present UcTCRp, a TCRβ-only framework for profiling MAIT- and iNKT-associated repertoire states in bulk immune repertoires. UcTCRp integrates V-gene context and CDR3β sequence features using a transformer-based representation pretrained on more than one million TCRβ sequences and supervised with curated cross-species MAIT, iNKT and conventional T cell references. The framework defines conserved model-informative TCRβ features, uses V-matched negative sampling to reduce germline-segment shortcuts, and generalizes across independent human and mouse datasets. In paired scRNA-seq/scTCR-seq datasets, UcTCRp recovered transcriptome-defined MAIT and iNKT cells and identified additional MAIT-like candidates supported by receptor evidence but missed by expression-only annotation. Bulk calibration against paired single-cell references and synthetic spike-in experiments established operating characteristics for repertoire-level abundance estimation. These results establish unpaired TCRβ repertoires as an actionable substrate for reconstructing unconventional T cell-associated immune states, enabling archived repertoire resources to be repurposed for systems-level studies of tissue immunity, disease and therapeutic response.

## Introduction

The human T cell receptor (TCR) repertoire exhibits extraordinary diversity, underpinning adaptive immunity across infections, tumors, autoimmunity, and immune-mediated tissue pathology^1^. Advances in immune repertoire sequencing have generated vast datasets that capture TCR diversity at unprecedented scale^2^. Bulk TCRβ sequencing, in particular, enables systematic characterization of clonal architecture, antigen-driven expansion, and repertoire diversity across large clinical and population cohorts^3^. However, most bulk repertoire datasets lack paired TCRα chains, transcriptomes, and phenotypic measurements. This limitation restricts the ability to assign observed clonotypes to defined T cell populations, especially rare or specialized lineages whose identities are typically established using information not directly encoded in bulk TCRβ data.

Among the cell populations most affected by this interpretability gap are unconventional αβ T cells, including mucosal-associated invariant T (MAIT) cells and invariant natural killer T (iNKT) cells^4^. MAIT cells recognize MR1-presented microbial riboflavin-derived metabolites^5^, whereas iNKT cells recognize CD1d-presented lipid antigens^6^. Although iNKT cells are typically rare in human peripheral blood and MAIT cell frequencies vary substantially across blood, age, disease state, and tissue context, both lineages are enriched or functionally specialized at barrier and non-lymphoid sites and can mount rapid effector responses^4^. Through cytokine production, cytotoxicity, and tissue-localized immune regulation, MAIT and iNKT cells contribute to antimicrobial defense^7,8^, tissue surveillance^9,10^, inflammation^11–13^, tumor immunity^14–16^, metabolic regulation^17,18^, and immune homeostasis^19^. Thus, the ability to recover MAIT- and iNKT-associated signals from existing TCRβ datasets would substantially expand opportunities to study unconventional T cell immunity across infection, aging, cancer, inflammatory disease, and tissue-specific immune perturbations. Yet current experimental approaches have not scaled to the breadth of available TCRβ sequencing resources.

Existing MAIT and iNKT annotation strategies rely on experimental or molecular information that is typically absent from bulk TCRβ datasets^20–22^. Antigen-loaded MR1 and CD1d tetramers provide direct cellular detection in targeted experiments^20,21^, whereas flow cytometric marker combinations define phenotypic subsets in fresh or well-preserved samples^23^. Single-cell RNA sequencing (scRNA-seq) coupled with paired V(D)J profiling further links receptor identity with transcriptional state^24^, and canonical TCRα-chain features, such as *TRAV1-2–TRAJ33/20/12* for human MAIT cells and *TRAV10–TRAJ18* for human iNKT cells^25^, are widely used as high-confidence molecular identifiers. Yet none of these measurements is available in most archived bulk TCRβ datasets, limiting their use for retrospective or population-scale studies of unconventional T cell immunity. Recent computational work has suggested that MAIT-associated receptor features can be learned from TCR sequence data, supporting the feasibility of TCR-based inference for unconventional T cell annotation^26^. Moreover, experimental evidence that the MAIT TCRβ chain contributes to ligand- and infection-dependent discrimination provides a biological basis for extracting lineage-relevant information from β-chain sequences^27^. It remains unclear, however, whether TCRβ-encoded lineage constraints extend beyond MAIT cells, generalize across species and can be calibrated into repertoire-level state estimates.

To address this gap, we present UcTCRp, a framework for inferring MAIT- and iNKT-associated immune states from TCRβ sequencing alone. UcTCRp captures conserved sequence features within unconventional T cell repertoires, integrates V-gene context and CDR3β composition, and converts single-clonotype predictions into calibrated repertoire-level estimates. We benchmarked UcTCRp across curated reference repertoires, independent cohorts, paired single-cell TCR and transcriptomic data, spike-in mixtures, and sequencing-depth perturbations. By enabling inference of MAIT- and iNKT-associated states from bulk and archival TCRβ datasets, UcTCRp provides a scalable and generalizable strategy to link TCR sequence architecture with unconventional T cell immunity, disease states, and potential therapeutic response.

## Results

### Conserved TCRβ features provide a basis for MAIT and iNKT inference

We first asked whether MAIT and iNKT TCRβ repertoires contain reproducible sequence information that could support TCRβ-only inference. To address this, we compiled a comprehensive dataset from the publicly accessible manually maintained database, UcTCRdb^28^. The dataset encompassed 20,117 MAIT TCRβs and 2,269 iNKT TCRβs from humans, as well as 1,786 MAIT TCRβs and 2,310 iNKT TCRβs from mice. Given the pivotal role of V and J genes in shaping the TCR, we first examined the gene usage patterns of these unconventional TCRβs compared to conventional CD4^+^ and CD8^+^ TCRβs. Despite the widely reported differences between conventional CD4^+^ and CD8^+^ T cells^29,30^, we found that *TRBV20-1, TRBV6-1,2,3,4, TRBV4-2,3 and TRBV15* were preferentially utilized in human MAIT cells, and *TRBV25-1* was significantly enriched in human iNKT cells compared with both CD4^+^ and CD8^+^ TCRβs (Figure 1A). In mice, *Trbv13-2,3, Trbv19 and Trbv4* were dominant in MAIT cells, while *Trbv13-1,2* and *Trbv29* were enriched in iNKT cells (Figure 1B). Across both human and mouse MAIT and iNKT cells, J gene usage was relatively diverse despite some subset-specific preferences (e.g., *TRBJ2-1,3* in human MAIT/iNKT and *Trbj2-3* in mouse MAIT cells), contrasting with the strong restriction and enrichment characteristic of V gene usage (Figure 1A, B). While these patterns are consistent with previous reports^31,32^ and reinforce the notion that MAIT and iNKT cells relay on highly restricted V segments linked to their evolutionarily conserved antigen specificity, our analysis of D gene reveals an additional layer of bias. Specifically, in the D gene annotation subsets of MAIT and iNKT TCRβ, we observed a statistically significant enrichment of the *TRBD2* gene fragment compared to conventional T cells (Figure 1C), while the *Trbd1* gene was significantly elevated in iNKT cells in the mouse dataset (Figure 1D). In addition to differences in gene usage, the unconventional TCRβs show a subtle but significant increase in CDR3 length compared to conventional T cells in both species (Figure 1E). Nevertheless, amino-acid conservation analysis shows that, despite biased gene usage and longer CDR3s, the CDR3 regions of unconventional TCRβ sequences remain highly diverse, as illustrated by the sequence-logo plots (Figure 1F, G).

**Figure 1.**
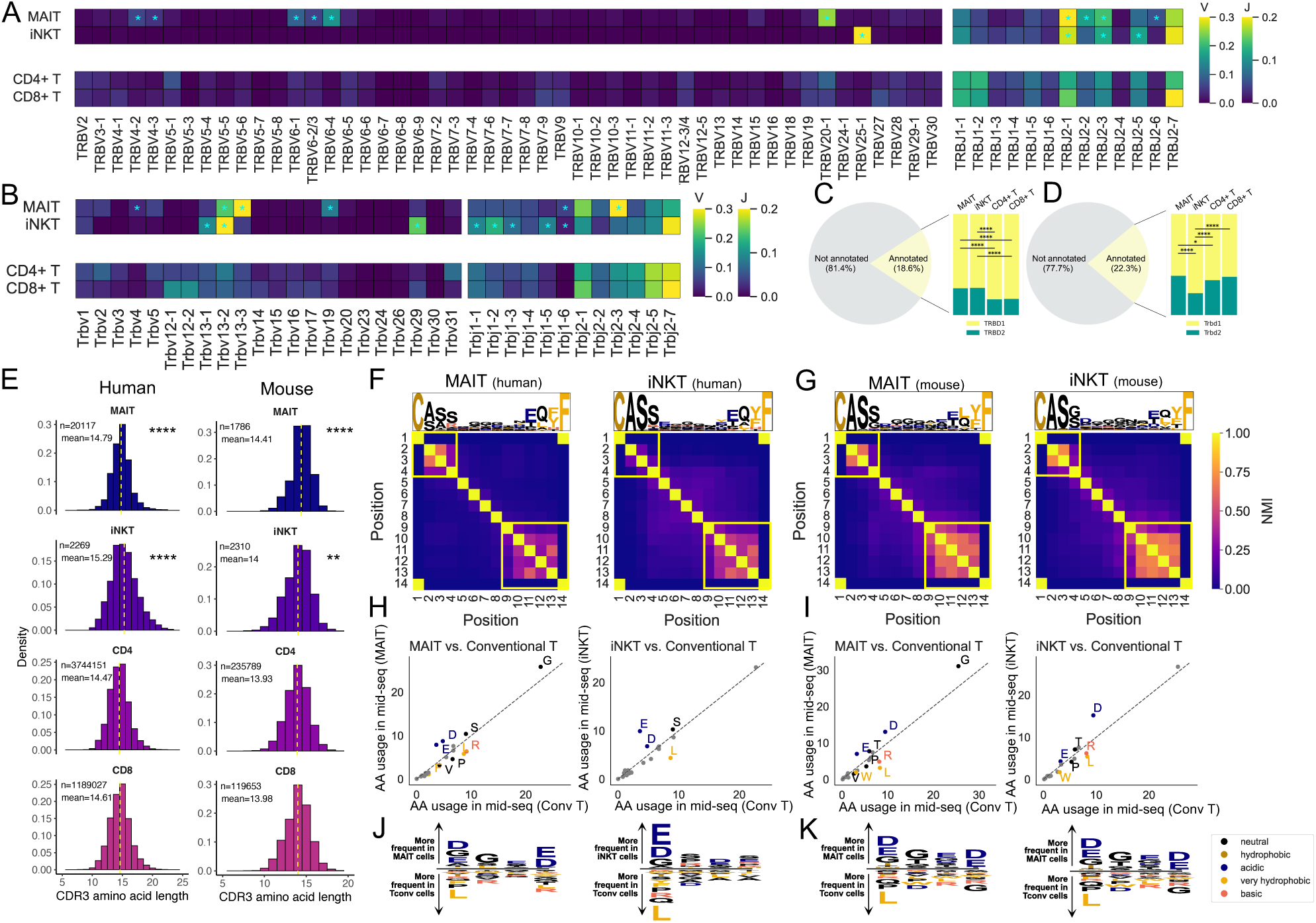
Cross-species reference repertoires define model-informative TCRβ features. **(A–B)** Heatmaps showing TRBV and TRBJ gene usage in MAIT and iNKT cells from human (A) and mouse (B), with conventional CD4⁺ and CD8⁺ T cells as controls. The color scale indicates the relative proportion of each gene’s usage. Asterisks (*) indicate significant gene-usage differences compared with conventional T cells (both CD4⁺ and CD8⁺), with FDR-adjusted p < 0.05 by Fisher’s exact test and an odds ratio > 1. **(C-D)** Proportional bar plots showing TRBD gene usage in a subset of human (C) and mouse (D) MAIT and iNKT cells, with conventional CD4⁺ and CD8⁺ T cells as controls. Asterisks (****) indicate FDR-adjusted p < 0.0001 by Fisher’s exact test. **(E)** Distributions of CDR3β amino acid lengths across unconventional and conventional T cells in human and mouse. Differences between MAIT, iNKT and conventional groups were assessed using Wilcoxon rank-sum tests with FDR adjustment (two-sided *p < 0.05; **p < 0.01; ***p < 0.001; ****p < 0.0001). **(F–G)** Sequence logos and position-wise normalized mutual information across the CDR3β region of fixed length (14 amino acids), highlighting germline-constrained (V- and J-encoded) versus highly recombined central regions. **(H–I)** Amino acid usage profiles in the central CDR3β “mid-seq” region among unconventional and conventional T cells. Only amino acids with significant differences by Wilcoxon rank-sum test after FDR adjustment (p < 0.05) and an absolute mean frequency difference greater than 1% are shown. **(J–K)** Position-specific enrichment of amino acid residues at positions 5–8 in CDR3β sequences of fixed length (14 amino acids) in human and mouse subsets.

To further dissect the compositional constraints underlying unconventional TCRβ sequences, we quantified normalized mutual information (NMI) across all residue pairs in CDR3β loops. In both human and murine samples, we observed elevated NMI at the N- and C-terminal regions of the CDR3β loop of fixed length (14 amino acids) (Figure 1F, G), as well as across other lengths (12, 13, 15 amino acids, Supplementary Figure 1), consistent with these positions being derived from germline-encoded V and J gene segments. In contrast, the central residues (positions 5-8 of length 14), which are shaped predominantly by random nucleotide additions during thymic recombination, exhibited markedly lower inter-residue dependence (Figure 1F, G). This pattern highlights somatic diversification of the unconventional CDR3β loops.

Given this contrast and in light of the conserved ligand recognition modes of unconventional αβ T cells^33–35^, we next asked whether the compositionally diverse CDR3β cores might nonetheless encode subset-specific features relevant to antigen engagement. We then compared amino acid usage between the central positions of unconventional and conventional CDR3βs. Strikingly, the biochemical composition of the unconventional CDR3β core was highly conserved between human and mouse. In both species, MAIT cells exhibited strong enrichment of glycine (G) together with negatively charged amino acids, specifically aspartic acid (D) and glutamic acid (E), whereas iNKT cells were consistently enriched for these negatively charged residues (D and E) (Figure 1H, I).

Although minor interspecies differences were also observed, for example, human MAIT and iNKT cells were enriched in serine (S), whereas their murine counterparts preferentially used threonine (T) (Figure 1H, I), this cross-species consistency suggests that these biochemical features in the CDR3β core may be functionally relevant for antigen recognition in each unconventional T cell subset. Motif analysis further revealed that such amino acid usage biases are broadly distributed across the central region of the CDR3β (Figure 1J, K), suggesting that although CDR3β regions are inherently diverse, unconventional T cells retain conserved amino acid preferences that likely underlie their capacity to engage invariant microbial or self-derived ligands.

To further substantiate these findings, we conducted additional analysis by subdividing conventional T cells into three distinct subtypes: CD4^+^ conventional T cells (Tconv), CD4^+^ regulatory T cells (Tregs), and CD8^+^ T cells. The results were mostly consistent with the initial observations (Supplementary Figure 2). Additionally, to address potential confounding effects, we refined our analysis to exclusively focus on TCRβs that shared the same V gene usage. Encouragingly, the identified amino acid usage patterns mostly persisted in both human (Supplementary Figure 3) and mice (Supplementary Figure 4), suggesting that CDR3β composition is shaped by selective pressures beyond germline-encoded V gene constraints. These collective findings underscore the consistency and prevalence of specific amino acid preferences or structural features within the intermediate sequences of unconventional T cells, emphasizing the potential functional significance of these patterns in the recognition of conserved antigens.

### UcTCRp integrates V-gene context and CDR3β features to classify unconventional TCRβs

Having established that MAIT and iNKT TCRβ repertoires contain conserved sequence constraints, we next developed UcTCRp, a TCRβ-only model designed to learn these constraints and classify unconventional TCRβ sequences. Considering the relatively limited fraction and diversity of unconventional TCRs (∼1-10% of the total T cell pool), we recognized the necessity of fortifying the model with robust stability and generalization capabilities. In recent years, the feature extraction capability of large-scale language models and their ability to capture underlying grammatical rules and hidden sequence patterns in a self-supervised manner based on unlabeled datasets have been demonstrated and successfully applied to protein sequence analysis^36,37^. Therefore, we construct a TCRβ multi-classification model by incorporating advanced pre-training technique based on TCR language model (Figure 2A).

**Figure 2.**
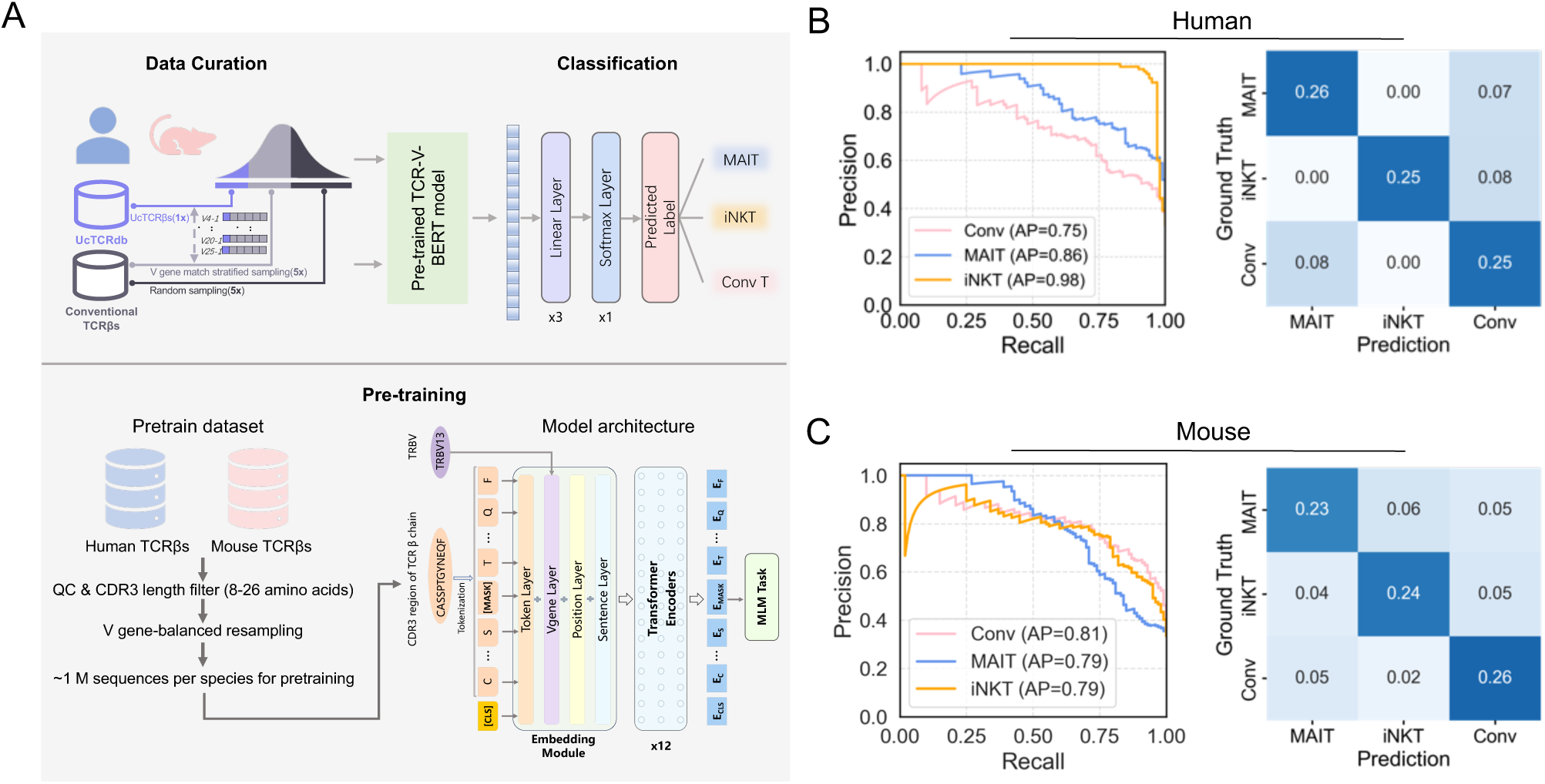
UcTCRp architecture and sequence-level classification performance. **(A)** Overview of the UcTCRp workflow. Input TCRβ sequences are encoded using V-gene context and CDR3β sequence, passed through a pretrained transformer representation, scored as MAIT-, iNKT- or conventional-associated states **(B, C)** Precision-recall curves and normalized confusion matrices illustrating model performance on human (B) and mouse (C) validation sets.

Specifically, to achieve a robust representation of TCRβs, we meticulously trained a Bidirectional Encoder Representation from Transformers (BERT)^38^ model on a dataset comprising one million TCRβ sequences at first step. This model, denoted as TCR-V-BERT, comprises 4 embedding layers and 12 transformer encoder modules, incorporating both V gene and CDR3 amino acid sequences as input. The J gene was excluded due to its limited variability and redundant information relative to the CDR3β region, which already captures its contribution to sequence composition.^39,40^ To capture the underlying syntax and semantics of TCRβ sequences, we employed a masked language modeling (MLM) pretraining strategy, which enables the model to learn generalizable sequence representations and global dependencies across V and CDR3 regions. The resulting embeddings provide informative features that can be transferred to downstream tasks, such as predicting antigen specificity or classifying T cell subsets. For our classification objective, we extracted sequence embeddings from the pretrained model and passed them through three fully connected layers trained with a weighted cross-entropy loss to address class imbalance in multi-label prediction.

During model training, we accounted for the known enrichment of specific V genes in unconventional TCRβs by implementing a sampling strategy to ensure balanced representation. For each unconventional TCRβ, 5-fold conventional TCRβs were randomly selected to match V gene usage and 5-fold were selected without V gene matching from publicly available repertoires (see Methods). This strategy simultaneously controlled for V gene biases and preserved realistic background variability, thereby mitigating the risk of the model learning V gene driven shortcuts. The resulting dataset, comprising both conventional and unconventional sequences, was randomly partitioned into training and testing sets at a 9:1 ratio. An independent validation set was constructed by sampling 100 sequences from each of the four defined subsets. Full details are summarized in Supplementary Tables 1 and 2.

As shown in Figure 2B, the human model demonstrated high classification performance across all unconventional T cell subsets, with area under the precision-recall curve (AUPRC) values of 0.86 for MAIT cells and 0.98 for iNKT cells. Most misclassifications occur between MAIT cells and conventional T cells rather than among unconventional subsets. This may reflect the presence of unannotated or latent MAIT TCRβs within the conventionally labeled negative samples, particularly given the relatively high prevalence of MAIT cells in peripheral blood and potential overlaps in V gene usage. The mouse model displayed slightly reduced performance relative to the human model, though it consistently achieved AUPRC values close to 0.8 across subsets (Figure 2C). Misclassification between murine iNKT and MAIT cells was more frequent, likely due to their shared V gene usage and convergent CDR3 features, as observed in Figure 1B, I and K.

### UcTCRp generalizes across independent cohorts and species

To further assess the robustness and generalizability of UcTCRp, we evaluated the model on three independent external cohorts that were not used during model development. These datasets included human MAIT TCRβ sequences from an independent study (n = 17,772)^41^, human MAIT (n = 17,340) and iNKT (n = 324) TCRβ sequences from a second cohort^42^, and mouse MAIT TCRβ sequences (n = 845)^43^ (Supplementary Table 3). UcTCRp retained strong performance across all validation datasets. The model achieved AUROC values of 0.87 in both human MAIT validation datasets, 0.92 for human iNKT cells and 0.85 for mouse MAIT cells (Figure 3A). Classification performance was also stable across additional metrics, with F1 scores of 0.81, 0.81, 0.86 and 0.76, respectively (Figure 3A). Thus, the pretrained model discriminated unconventional TCRβ sequences across independent human cohorts and retained predictive performance in mouse MAIT repertoires without retraining.

**Figure 3.**
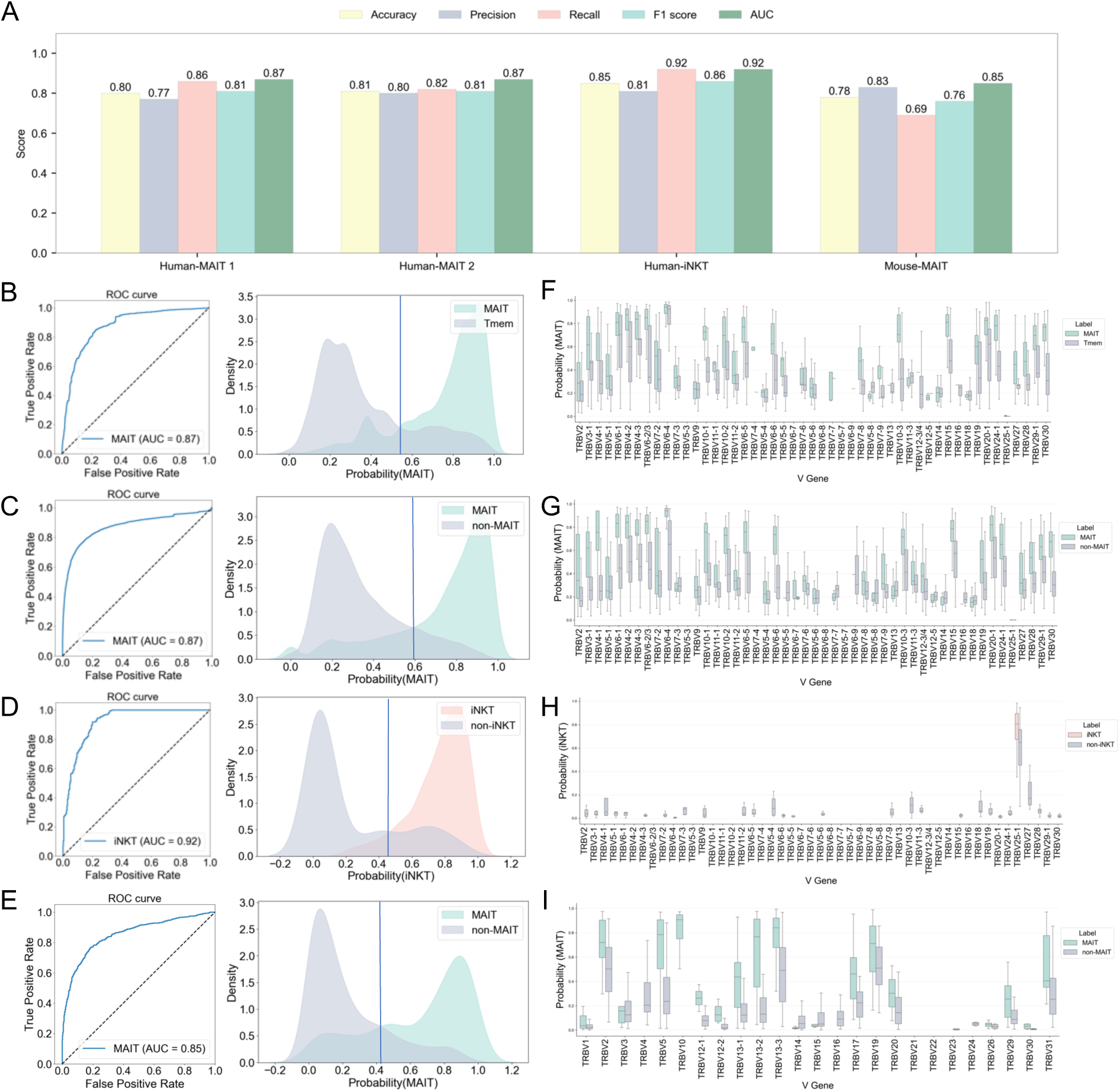
External validation of UcTCRp across independent datasets. **(A)** Model performance across four external validation settings, summarized by AUROC, accuracy, precision, recall and F1 score. **(B–E)** ROC curve and distributions of model prediction scores for positive and negative TCRβ sequences across four external validation cohorts: (B) human MAIT vs. non-MAIT memory T cells (Validation Set 1), (C) human MAIT vs. non-MAIT conventional T cells (Validation Set 2), (D) human iNKT vs. non-iNKT conventional T cells, and (E) mouse MAIT vs. non-MAIT conventional T cells. **(F–I)** Score distributions stratified by V gene usage within each dataset (corresponding to B–E), showing that separation between positive and negative sequences persists across diverse V gene backgrounds.

Prediction score distributions confirmed these results. In both human MAIT validation datasets, MAIT TCRβ sequences showed a clear shift toward higher predicted MAIT probabilities relative to non-MAIT memory or conventional T cells (Figure 3B, C). Human iNKT sequences were similarly enriched at high predicted iNKT probabilities compared with negative controls (Figure 3D). Mouse MAIT sequences also showed higher predicted MAIT scores than non-MAIT conventional T cells, although separation was more moderate than in the human datasets (Figure 3E).

We next stratified prediction scores by V gene usage to determine whether classification was explained solely by germline gene enrichment. In the human MAIT datasets, high predicted scores were enriched in MAIT-associated Vβ backgrounds, including *TRBV20-1* and *TRBV6-4*, but MAIT sequences remained separated from negative sequences across multiple V genes (Figure 3F, G). In the human iNKT dataset, high scores were concentrated in *TRBV25-1*, consistent with known iNKT-associated Vβ usage (Figure 3H). Mouse MAIT sequences also showed V gene-stratified separation despite greater variability across strata (Figure 3I). These results confirm that the model captures both CDR3-intrinsic features and biologically relevant germline context, enabling accurate discrimination of unconventional TCRβs across species.

### Single cell validation supports β-chain-based unconventional T cell annotation

Having established generalization across independent datasets and species, we next evaluated UcTCRp using orthogonal single-cell evidence. We applied the model to a large public scRNA-seq dataset^44^ coupled with paired V(D)J profiling to compare β-chain-based predictions with transcriptomic annotations and canonical α-chain-based MAIT definitions. The dataset contained 1.1 million T cells, including 483,838 cells with paired transcriptome and TCR information across 35 annotated cell types (Figure 4A). The original study annotated two MAIT-related groups: MAIT cells (n = 48,158; 9.95%) and CD8.c16.MAIT.SLC4A10 cells (n = 2,990; 0.62%). iNKT cells were not annotated as a separate population in the original dataset. UcTCRp scores were computed for 431,515 cells with suitable TCRβ information. We then compared three MAIT annotation layers: original transcriptomic annotation, canonical α-chain annotation based on *TRAV1-2* paired with *TRAJ33*, *TRAJ20* or *TRAJ12*, and β-chain annotation based on UcTCRp scores.

**Figure 4.**
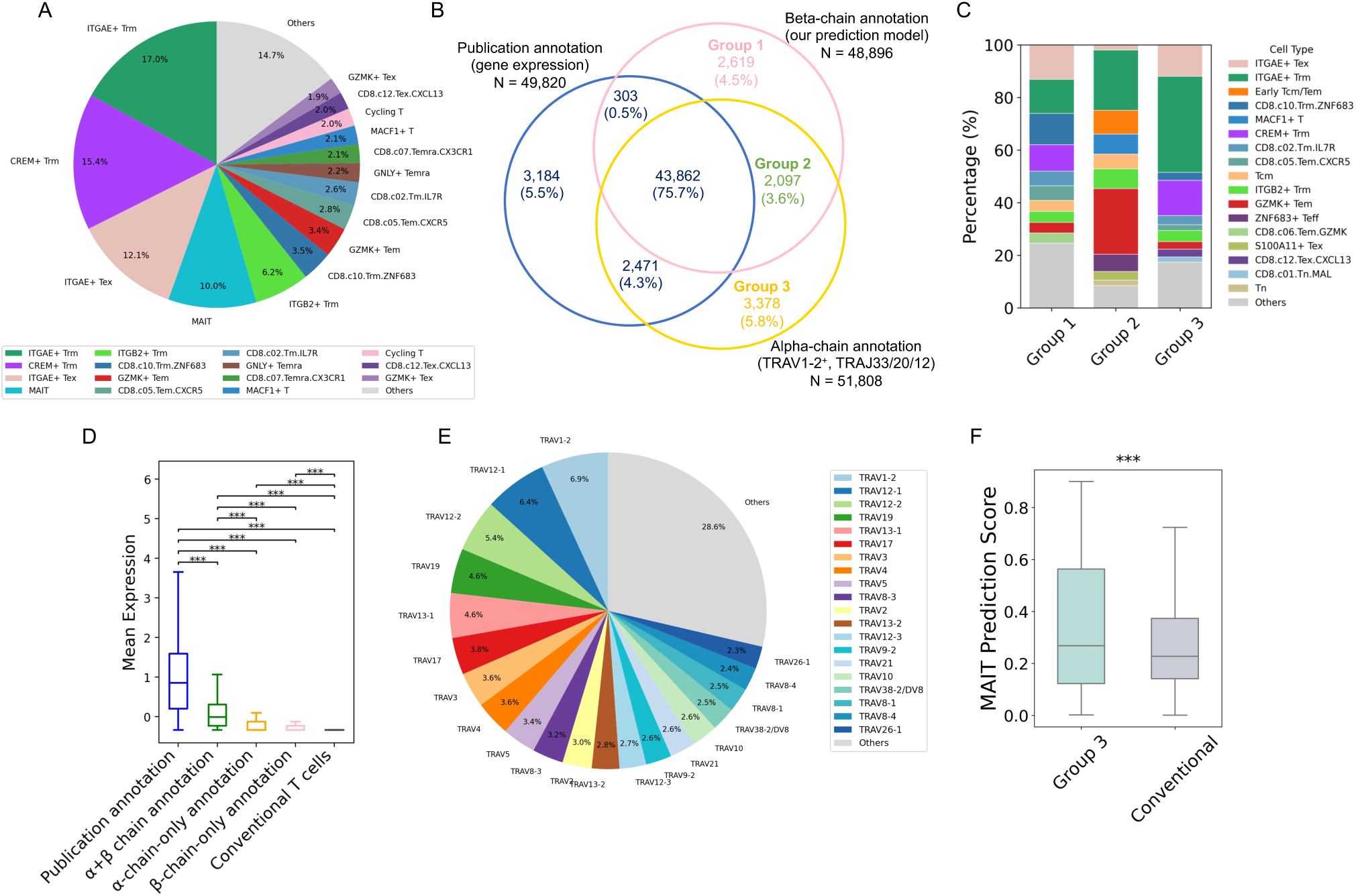
Single-cell validation using paired scRNA-seq and scTCR-seq data. **(A)** Cell type proportions in the original dataset (n=483,838). Only the top 15 most abundant cell types are shown, with the remaining types grouped into “Others”. **(B)** Overlap between MAIT cells defined by β-chain prediction score and α-chain annotation, and MAIT cells with original annotation in the publication. Group1: β-chain-only annotation (n=2,619); Group2: α+β chain annotation (n=2,097); Group3: α-chain-only annotation (n=3,378). **(C)** Distribution of original transcriptome-defined cell types among Groups 1–3. The ten most abundant cell types in each group are shown; remaining cell types are grouped as “Others”. **(D)** Expression of MAIT cell gene markers (*ZBTB16*, *KLRB1*, *SLC4A10*) across five cell groups: original transcriptome-annotated MAIT cells, α+β-supported cells outside the original MAIT label, α-chain-only cells outside the original MAIT label, β-chain-only cells outside the original MAIT label and conventional T cells. Values indicate mean marker expression. Pairwise comparisons were performed using two-sided Mann–Whitney U tests with Bonferroni correction. Significant differences are indicated as: *, P < 0.05; **, P < 0.01; ***, P < 0.001. **(E)** TCRα V-gene usage among cells annotated as MAIT by UcTCRp β-chain prediction but not by canonical MAIT α-chain criteria. **(F)** Distribution of UcTCRp MAIT prediction scores in α-chain-only cells outside the original MAIT label and conventional T cells. Statistical significance was assessed using a two-sided Mann–Whitney U test.

The three annotation strategies showed substantial but incomplete overlap (Figure 4B). In total, 43,862 cells were jointly supported by transcriptomic, α-chain and β-chain annotations, corresponding to 75.7% of the combined MAIT-like set. UcTCRp and α-chain annotation also identified additional cells not labeled as MAIT in the original transcriptomic annotation. To characterize these discordant cells, we defined three groups: β-chain-only annotation (Group 1; n = 2,619), α+β chain annotation outside the original MAIT label (Group 2; n = 2,097), and α-chain-only annotation outside the original MAIT label (Group 3; n = 3,378) (Figure 4B). These cells were distributed across multiple transcriptomic cell types and were relatively enriched among ITGAE+ tissue-resident memory-like cells (Figure 4C), suggesting that gene-expression-based annotation alone may miss MAIT-like cells in certain phenotypic states.

Marker expression supported this interpretation. Original transcriptomic MAIT cells showed the highest expression of MAIT-associated markers including *ZBTB16*, *KLRB1* and *SLC4A10*. Among cells not originally labeled as MAIT, the α+β-supported group showed the strongest marker expression, whereas α-only and β-only groups also showed significantly higher MAIT marker expression than conventional T cells (Figure 4D). These results indicate that TCR-based annotations capture biologically relevant MAIT-like cells that may be undercalled by transcriptomic clustering alone.

We further examined discordance between α- and β-chain evidence. Cells annotated by β-chain score alone tended to pair with non-classical MAIT-like α chains rather than the canonical *TRAV1-2–TRAJ33/20/12* combination (Figure 4E), suggesting that UcTCRp may help recover MAIT-like cells with non-classical α-chain usage. Conversely, α-chain-only cells that did not pass the UcTCRp threshold nevertheless had significantly higher β-chain prediction scores than conventional T cells (Figure 4F), indicating that α- and β-chain MAIT features are partially coupled even when one chain falls below the annotation threshold. Moreover, UcTCRp detected a small group of transcriptome-labeled MAIT cells with iNKT-like β-chain scores. Most of these cells carried *TRAV10–TRAJ18* α chains, providing independent receptor-level support for this assignment. Although canonical *TRAV10–TRAJ18/TRBV25-1* cells were rare in this dataset (n = 97), this concordance suggests that UcTCRp can recover rare iNKT-like signals not resolved by the original transcriptomic annotation.

### Aggregated UcTCRp scores estimate unconventional T cell states in bulk repertoires

Having established that UcTCRp accurately classifies unconventional TCRβ sequences, we next tested whether sequence-level scores could be aggregated to estimate unconventional T cell states in bulk repertoires. We therefore evaluated sample-level calibration using a scRNA-seq dataset from 305 peripheral blood samples of 166 healthy donors (Validation Set 2)^42^, with a median of 2,545 cells per sample having paired TCR sequences and transcriptome-defined cell type annotations. For each sample, UcTCRp scores were computed from available TCRβ sequences and aggregated as the fraction of sequences exceeding a series of decision cutoffs ranging from 0.5 to 0.9. These estimates were then compared with transcriptome-defined MAIT and iNKT cell abundances to assess whether β-chain-based predictions captured sample-to-sample variation in unconventional T cell states.

Aggregated UcTCRp estimates tracked measured MAIT abundance across all tested cutoffs, with correlations increasing as the cutoff became more stringent. Spearman correlations ranged from ρ =

0.291 at a cutoff of 0.5 to ρ = 0.672 at a cutoff of 0.9, with all comparisons statistically significant (Figure 5A). For iNKT cells, estimated frequencies also showed significant positive correlations across all cutoffs, with correlations peaking at 0.8 (ρ = 0.510, *P* = 5.1 × 10^-8^) and remaining significant at 0.9 (ρ = 0.371, *P* = 1.3 × 10^-4^; Figure 5B). Because downstream bulk analyses prioritize high-confidence subset-associated signals over maximal sensitivity in a single calibration dataset, we used a unified conservative cutoff of 0.9 for both MAIT and iNKT estimates. To further test quantitative recovery in controlled bulk-like settings, we then generated simulated datasets by spiking unconventional TCRβ sequences from the validation set into unseen conventional CD4⁺ or CD8⁺ TCRβ backgrounds at varying proportions. Across all simulations, model-predicted frequencies exhibited strong positive correlations with true input values for each unconventional subset, in both CD4⁺ and CD8⁺ compartments (Figure 6A, B), confirming the model’s ability to resolve subset-level abundance in complex repertoires.

**Figure 5.**
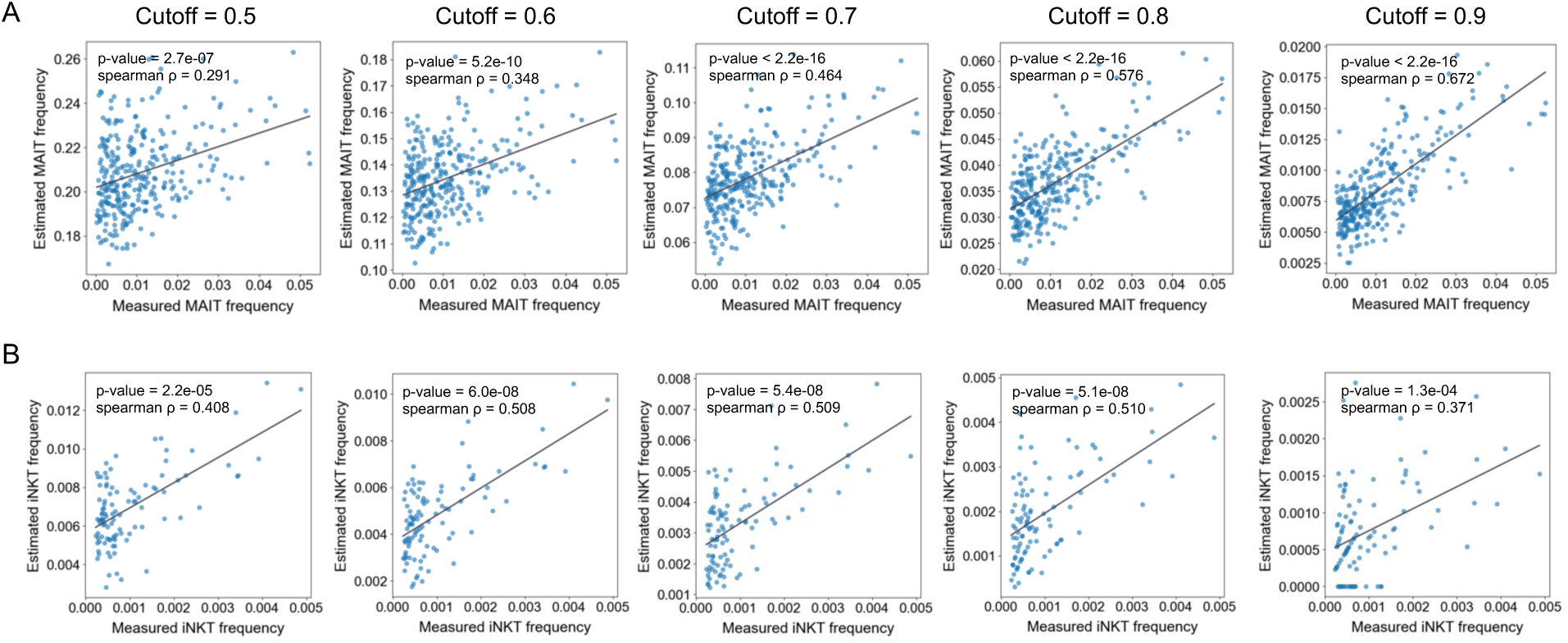
Sample-level calibration of UcTCRp estimates across score cutoffs. For each sample in Validation Set 2, MAIT and iNKT frequencies were estimated as the fraction of TCRβ sequences exceeding UcTCRp score cutoffs from 0.5 to 0.9. Predicted frequencies were compared with transcriptome-defined MAIT and iNKT cell abundances **(A)** Correlation between predicted and true MAIT frequencies based on the number of TCRβ clonotypes. **(B)** Correlation between predicted and true iNKT frequencies based on the number of TCRβ clonotypes. Spearman correlation coefficients (r) and P values are indicated for each threshold.

**Figure 6.**
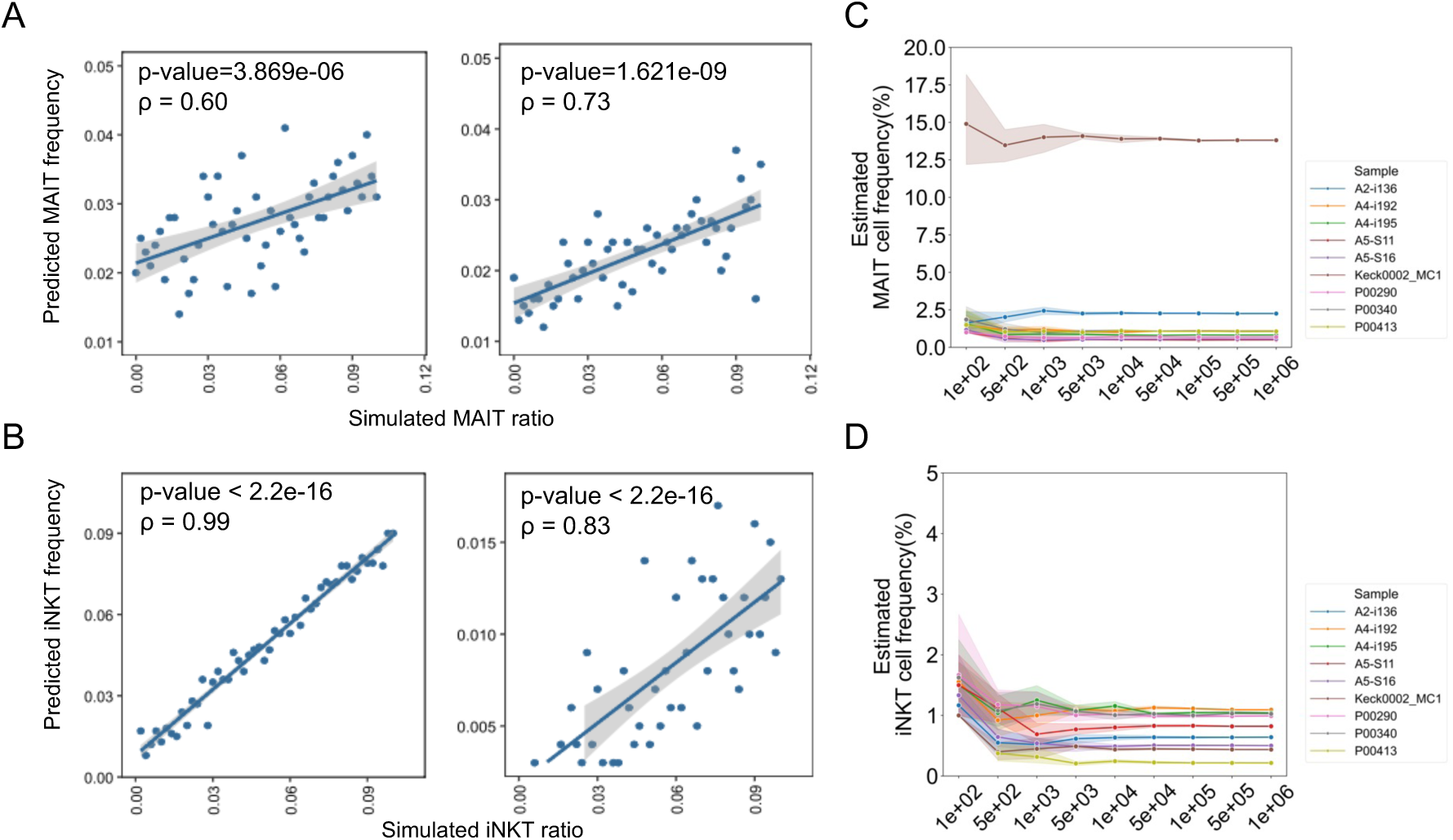
Quantitative recovery and sequencing-depth analysis of UcTCRp-based abundance estimates. **(A, B)** Spike-in simulations for MAIT (A) and iNKT (B) abundance estimation. Unconventional TCRβ sequences were mixed into unseen conventional CD4+ or CD8+ TCRβ backgrounds at defined proportions. The x-axis shows the true spike-in proportion, and the y-axis shows the UcTCRp-estimated proportion using a fixed score cutoff of 0.9. Spearman correlation coefficients and P values are shown for each simulated setting. **(C, D)** Downsampling analysis for MAIT (C) and iNKT (D) abundance estimation. Simulated repertoires were generated by subsampling TCRβ sequences from nine randomly selected donors across depths ranging from 100 to 1,000,000 unique TCRβ sequences. Predictions were performed using a fixed score cutoff of 0.9. Solid lines indicate mean predicted abundances across replicates, and shaded regions indicate 95% confidence intervals. The dashed line indicates the recommended minimum sequencing depth of 1,000 unique TCRβ sequences for stable repertoire-level estimation.

To assess the robustness of abundance estimation under varying sequencing depths, we further performed a downsampling analysis using bulk TCRβ repertoires from nine randomly selected healthy donors in the Emerson et al. cohort^45^. For each repertoire, TCRβ sequences were randomly subsampled to total sizes of 100 to 1,000,000. At each depth, subsampling was repeated ten times to control for stochastic variation, and model-based predictions of MAIT and iNKT cell frequencies were recorded. For each sampling depth, we computed the mean and variance of predicted frequencies across replicates. Predicted abundances of unconventional T cells remained stable when repertoire sizes exceeded 1,000 unique TCRβ sequences; however, estimates became highly variable below this threshold across all subsets (Figure 6C, D). Accordingly, we defined 1,000 unique TCRβs as the minimum input size required for reliable abundance estimation.

Together, these results establish a robust framework for inferring unconventional T cell subset abundances from bulk TCRβ sequencing, enabling scalable immunophenotyping across diverse samples and experimental conditions.

## Discussion

T cell immunity has traditionally been interpreted through the recognition of peptide antigens presented by MHC class I or class II molecules. This paradigm, however, does not fully capture unconventional, non-MHC-restricted T cells. MAIT and iNKT cells recognize conserved microbial or self-derived antigens presented by MR1 and CD1d, respectively, and contribute to tissue surveillance, antimicrobial defense and immune regulation^4^. Yet these lineages remain difficult to study at scale because their conventional identification relies on paired TCRα chains, antigen-loaded tetramers or cellular phenotypes, whereas most public and clinical immune-repertoire datasets contain only unpaired TCRβ sequences. Here we show that UcTCRp can extract learnable and calibratable MAIT- and iNKT-associated signals from such incomplete repertoire data, enabling single-cell annotation support and bulk repertoire-level inference without redefining these lineages by β chain alone.

A key implication of this work is that MAIT- and iNKT-associated β chains are not merely passive partners of invariant or semi-invariant α chains. Across human and mouse repertoires, these lineages showed recurrent TRBV and TRBJ biases, CDR3β-associated sequence features and conserved amino acid enrichment within the junctional core. These patterns were maintained across species and phenotypic contexts, suggesting that unconventional TCRβ repertoires are shaped by conserved selective pressures rather than by dataset-specific sampling effects. The persistence of predictive signal after V-matched negative sampling and V-stratified evaluation further indicates that these features are not reducible to germline-segment enrichment alone. Instead, MAIT and iNKT TCRβ repertoires appear to encode distributed lineage-associated constraints that can be learned from large-scale sequence data and calibrated for repertoire-level analysis.

The enrichment of negatively charged residues within the CDR3β core provides a plausible link between sequence-level constraints and antigen-recognition biology. Aspartic acid and glutamic acid residues may shape the electrostatic environment of the TCR–ligand interface or stabilize conformations compatible with MR1- or CD1d-restricted recognition. Structural studies of MAIT TCR–MR1–antigen complexes indicate that CDR3β can contribute to the TCR–MR1–antigen interface and modulate complex stability^33,34^, although direct antigen contacts by CDR3β are not consistently observed. Rather than acting as a deterministic contact motif, CDR3β may support recognition through conformational adaptability and induced-fit-like rearrangements, as illustrated by the conformational shift of E99β in the MR1–Ac-6-FP complex^34^. Similar principles have been described for iNKT TCR recognition of CD1d–lipid complexes^31^, where β-chain features can influence docking geometry and ligand discrimination. Thus, the conserved charge distributions identified here may reflect structural constraints that tune receptor engagement rather than simple sequence motifs. Future structure-aware models could test this hypothesis more directly.

These observations motivated the development of UcTCRp as a framework for extracting unconventional T cell-associated information from TCRβ-only data. Prior computational work has suggested that MAIT-associated receptor features can be learned from TCR sequence data^26^.

UcTCRp extends this concept by modeling multiple unconventional T cell lineages, incorporating human and mouse repertoires, and translating sequence-level predictions into repertoire-level estimates. Its design combines V-gene context, CDR3β sequence information and self-supervised transformer pretraining, allowing the model to learn distributed receptor features rather than rely on single-gene enrichment or simple motif rules. This architecture is aligned with the structure of most public and clinical repertoire resources, which often contain only TCRβ sequences.

The paired single-cell validation places this β-chain signal in the context of established annotation modalities. UcTCRp recovered transcriptome-defined MAIT cells and identified additional candidate MAIT-like cells supported by αβ receptor evidence but not captured by expression-only annotation. This is consistent with the expectation that transcriptomic clustering, canonical α-chain usage and β-chain inference capture overlapping but non-identical aspects of MAIT identity^41,46^. Transcriptional profiles vary with activation state, tissue localization and stimulation context^41,46^, whereas canonical α-chain rules are highly specific for classical MAIT cells but may miss non-classical MR1-reactive or MAIT-like receptor configurations^47,48^. In this setting, UcTCRp provides an orthogonal receptor-based annotation layer for paired single-cell datasets and a scalable inference layer for bulk datasets in which α-chain and transcriptomic information are absent.

At the repertoire level, aggregated UcTCRp scores enable MAIT- and iNKT-associated states to be approximated from bulk TCRβ data^49^. This is the setting in which the method has its greatest practical value. For large archived cohorts, the relevant question is often not whether a single clonotype can be definitively assigned to a lineage, but whether a sample carries increased or decreased unconventional T cell-associated receptor signal. By calibrating sequence-level scores against paired single-cell references and controlled spike-in mixtures, UcTCRp translates clonotype-level predictions into sample-level repertoire summaries (Figure 5, 6). These estimates should be interpreted as β-chain-associated lineage-state signals rather than exact flow-cytometric frequencies or proof of MR1/CD1d restriction. Within this boundary, the framework enables retrospective analysis of MAIT- and iNKT-associated variation across infection, aging, cancer, inflammatory disease and therapeutic response.

Nonetheless, several limitations remain. The biological interpretation of MAIT- and iNKT-associated CDR3β features remains partly inferential because TCR recognition is shaped by paired αβ receptor geometry, antigen-presenting molecule, ligand context and cellular state. Although UcTCRp detects lineage-associated receptor signals from TCRβ sequences, it does not establish antigen specificity, MR1 or CD1d restriction, activation state or functional subset identity for individual clonotypes.

Further validation through structural modeling, ligand-binding assays and antigen-loaded tetramer experiments will therefore be needed to determine whether recurrent β-chain features contribute directly to antigen recognition or instead reflect lineage-associated receptor constraints. Explicit modeling of antigen context, including microbial metabolite, lipid antigen and presenting molecule information, may further distinguish lineage-associated signals from ligand-specific recognition features. Finally, integrating orthogonal modalities such as transcriptomics, surface proteomics and antigen-reactivity measurements could enable more comprehensive models of unconventional T cell identity, state and function. Extending this framework to additional semi-invariant unconventional populations, including other MR1-reactive, CD1-restricted and HLA-E-restricted T cells^4^, will help determine whether partial receptor sequences can more broadly support inference of non-peptide and non-classical antigen recognition.

In summary, UcTCRp demonstrates that unpaired TCRβ repertoires encode structured, lineage-associated information about MAIT and iNKT immune states. By learning these constraints, validating them against single-cell and cross-species evidence, and calibrating them for bulk deployment, the framework links receptor sequence architecture to unconventional T cell biology at population scale. More broadly, this work supports a general principle for immune-repertoire analysis: incomplete receptor measurements can contain biologically organized cell-state information that, when appropriately modeled and calibrated, can transform archived clonotype catalogues into quantitative maps of immune-state variation.

## Materials and methods

### TCRβ Sequence Collection and Annotation

TCRβ sequences representing MAIT cells, iNKT cells, and conventional αβ T cells were compiled from multiple sources, including the UcTCRdb^28^ (a manually curated database of unconventional T cells) and primary dataset retrieved from published bulk TCR sequencing study^50^ with CD8⁺, CD4⁺, and Treg subsets. For inclusion, datasets were required to contain annotated CDR3β amino acid sequences with matched V and J gene assignments. The final dataset encompassed 22,386 unconventional TCRβ sequences from human samples and 4,096 from murine sources. Subsets included 20,117 MAIT and 2,269 iNKT TCRβs from human repertoires, and 1,786 MAIT and 2,310 iNKT TCRβs from murine repertoires. For conventional αβ T cells (CD4⁺, CD8⁺, and Treg), each subset comprised more than one million TCRβ sequences, serving as the negative class during model training and evaluation.

Rigorous quality control criteria were applied prior to downstream analysis. Only productive, in-frame CDR3β sequences were retained. Sequences were required to conform to IMGT-defined structural boundaries: beginning with a cysteine (C) and terminating in a phenylalanine (F), with lengths constrained to 8–26 amino acids. Sequences containing stop codons, nonstandard amino acid characters, or unresolved annotations were excluded. Duplicate entries were removed based on the combination of CDR3β amino acid sequence, V gene, and J gene identifiers. Given the heterogeneity in V and J gene annotations across datasets (e.g., differences between “TRBV” and “TCRBV” nomenclature, and inconsistent allele-level resolution), a unified gene-mapping dictionary was constructed based on the IMGT and Emerson et al.^45^ reference tables. In cases where multiple alleles were ambiguously assigned or reported as merged variants (e.g., “TRBV06-02/06-03”), a standardized composite label (“TCRBV06-02/03”) was assigned. V and J gene annotations not found in the curated list or corresponding to pseudogenes were excluded to ensure consistency in downstream embedding and classification tasks. For datasets lacking allele resolution or reporting ambiguous V gene calls (e.g., “TCRBV06-02,TCRBV06-03”), consensus-based collapsing rules were applied, and a majority-vote identifier was retained.

### Gene Usage and CDR3 Composition Analysis

To explore potential V gene usage preferences in unconventional T cells, the frequency of each V gene usage in each subtype were calculated and plotted on a heatmap. For compositional analysis, CDR3β sequences were grouped by length, and analyses were performed independently within each length category (e.g., L = 13, 14, 15, 16 amino acids). Within each group, normalized mutual information (NMI) was computed between all positional pairs to quantify the dependency of amino acid usage across the CDR3β sequence. NMI were calculated as following:

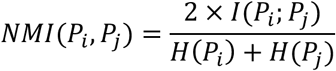

where *P_i_* denotes the amino acid at the i-th position within the CDR3 region, *P_j_* indicates any other amino acid at position denoted by j, *i, j* ∈ [1, *l*]. H(.) stands for cross entropy and I(X;Y) stands for mutual information, which is used to measure the relevance of pairwise amino acids:

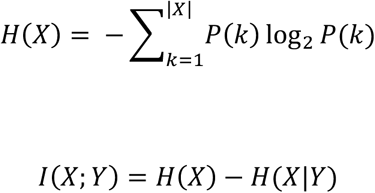

“Mid-seq” region was defined as follows: the first four and the last six residues of the CDR3 sequence were uniformly removed, and the intermediate sequences of CDR3 was retained as the middle sequence. We analyzed the amino acid usage of intermediate (“mid-seq”) positions of CDR3 sequences with lengths greater than 10 amino acids. Firstly, the frequency of 20 amino acids in the intermediate sequence of each CDR3 was calculated. The mean frequency of each amino acid across sequences was then calculated to obtain the average amino acid usage in each subtype of T cells. The frequency of each amino acid in MAIT and iNKT cells was plotted on a scatter plot in comparison with conventional T cells, in which those with no significant difference were colored in gray, and amino acids that deviated from the diagonal were highlighted with corresponding colors according to physicochemical properties (Figure 1H, I, Supplementary Figure 2). The above calculations were consistent when V gene restriction was added for analysis (Supplementary Figure 3, 4). In addition, sequences with a fixed CDR3β length of 14 amino acids were selected to generate sequence logo plots, enabling direct visual comparison of residue preference patterns between unconventional and conventional T cell subsets. Different amino acids were colored in accordance with physiochemical and biological properties.

### Transformer-Based Pretraining of TCR Embeddings (TCR-V-BERT)

To extract generalizable representations of TCRβ sequences and capture their intrinsic sequence grammar, we developed a TCR-specific transformer model (TCR-V-BERT) based on the BERT architecture. This model was pretrained on a large-scale corpus of unlabeled TCRβ sequences using a masked language modeling (MLM) objective, enabling downstream transfer to classification tasks involving unconventional T cell identification.

#### (1) Pretraining Dataset and Preprocessing

A total of 153,170,151 human TCRβ sequences annotated with V gene information were obtained from a publicly available cytomegalovirus (CMV)-associated repertoire study^45^. To ensure high-quality input, sequences containing illegal characters, stop codons, or invalid amino acid residues were removed. CDR3β sequences were required to be in-frame, productive, and within the length range of 8 to 26 amino acids. Duplicated entries were collapsed. Given the class imbalance across different TRBV gene families, we implemented a stratified downsampling strategy to ensure adequate representation of all V gene classes. Sequences associated with low-frequency V genes were fully retained, whereas those from high-frequency V genes were randomly subsampled, yielding a final balanced corpus of ∼1 million sequences for model pretraining. This dataset was split into training and validation sets at a 1000:1 ratio.

#### (2) Model Architecture

The TCR-V-BERT model adopts the standard BERT^38^ encoder structure. Inputs to the model consisted of tokenized CDR3β amino acid sequences and V gene identifiers. Each sequence was augmented with special tokens ([CLS]) and padded to a uniform length. Input representations were constructed from four types of embeddings:

- Token embeddings: 20 canonical amino acids (plus special tokens) mapped to 768-dimensional vectors.
- V gene embeddings: Unique V gene identifiers encoded and projected to 768 dimensions.
- Positional embeddings: Encoded token positions within the sequence to retain order information.
- Segment embeddings: Binary indicators denoting sequence type (i.e., TCRα versus TCRβ), following the standard BERT architecture. In the current study, all input sequences correspond to TCRβ chains, and this component is retained to support future extensions involving paired-chain (TCRαβ) modeling.

These embeddings were summed and passed through a 12-layer transformer encoder, each layer comprising multi-head self-attention and feed-forward submodules with layer normalization.

#### (3) Pretraining Objective and Optimization

We employed the MLM task, wherein 15% of amino acids in each input sequence were randomly masked and the model was trained to predict the masked residues. The MLM head consisted of a dense projection layer and softmax classifier over the amino acid vocabulary. The objective function was a weighted cross-entropy loss:

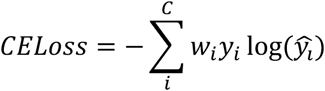

where *w_i_* is the weight of class i, *y_i_* is the true label of class i, *ŷ_i_*) is the predicted label of class i, and C represents the total number of classes. Model training was implemented in PyTorch v1.12.1 with the Adam optimizer (learning rate = 3×10⁻⁵, betas = 0.9 and 0.999), batch size = 64, and GELU activation. Training was run for 50 epochs on a Supermicro 7049GP-TRT server equipped with NVIDIA GeForce RTX 2080Ti GPUs under Ubuntu 18.04. All training and evaluation pipelines were orchestrated using Python 3.9.3 and JupyterLab 2.1.5.

#### (4) Cross-Species Extension

To support downstream analyses in murine TCR repertoires, we constructed an additional TCR-V-BERT model pretrained on mouse TCRβ sequences following the same preprocessing, downsampling, and architectural pipeline. This parallel model enabled unbiased embedding and classification of both human and mouse TCRs across all experiments.

### Supervised Classification of Unconventional TCRs (UcTCRp)

To enable automated identification of MAIT and iNKT TCRβ sequences from bulk repertoires, we fine-tuned the pretrained TCR-V-BERT embeddings using a supervised multi-class classifier. The task was formulated as a three-class classification problem: MAIT, iNKT, and conventional TCRβs.

#### (1) Classifier Architecture

The classification module (UcTCR-Predictor) was implemented as a three-laye feed-forward neural network. Input embeddings were computed as the mean of hidden states from the 8^th^ transformer layer of the 12-layer TCR-V-BERT model, a strategy empirically shown to improve generalization in biological sequence modeling tasks^37^. The network consisted of two fully connected layers (from 768 to 256 and from 256 to 64 dimensions), each followed by batch normalization and a ReLU activation. Dropout was applied after each hidden layer (rates = 0.3), and a final linear layer mapped the 64-dimensional representation to three output logits. Class probabilities were obtained by applying a softmax function to these logits. The model was trained using a weighted cross-entropy loss to address class imbalance, with class weights set inversely proportional to the observed class frequencies. Because human iNKT cells predominantly utilize TRBV25-1, we introduced an additional training label for conventional TRBV25-1⁺ TCRβ sequences to prevent the model from overfitting to this feature; at inference, predictions for this auxiliary class were merged with those for the conventional T-cell class.

#### (2) Training and Evaluation Dataset Construction

Training and validation sets were assembled from curated high-confidence human and murine TCRβ sequences with known subset identities (see above). To mitigate biases due to unequal class representation and TRBV usage imbalance, we adopted a dual-sampling strategy for generating negative (conventional) TCRs:

- V gene–matched negatives (×5): For each unconventional TCR, five conventional sequences were sampled that shared the same TRBV gene.
- Unmatched negatives (×5): Five additional conventional sequences were randomly sampled from unrelated donors to capture global repertoire diversity.

All sequences were required to be productive, in-frame, and contain valid TRBV, TRBJ, and CDR3β annotations. To prevent label leakage, conventional sequences with similar V gene and Hamming distance <3 to any unconventional CDR3β were excluded. The final training dataset included 243,438 human and 42,823 mouse sequences. Data were randomly split into training (90%) and test (10%) sets. A separate external validation cohort was constructed by independently sampling 100 sequences per class.

#### (3) Model Training and Performance Evaluation

Fine-tuning was performed using the Adam optimizer with L2 regularization (weight decay=2×10⁻^6^, learning rate = 8×10⁻^5^, batch size = 256) for 16 epochs. Early stopping was applied based on test loss. Classification performance was evaluated using AUROC, precision, recall, and F1 score, computed separately for each class.

#### (4) External Validation

To evaluate the generalizability of our model, we tested its performance on three independent external datasets encompassing different platforms, cell types, and species:

> Validation Set 1: scTCR-seq of human MAIT and non-MAIT memory T cells (n = 17,772), with subset identity determined by antigen stimulation and tetramer enrichment.^41^
>
> Validation Set 2 and 3: scRNA-seq and scTCR-seq of human PBMCs with annotated MAIT (n = 17,340) and iNKT (n = 324) cells, defined based on gene expression–derived cluster identities.^42^
>
> Validation Set 4: TCRβ repertoires from flow-sorted MAIT T cells isolated from mouse thymus (n = 845), identified via MR1 tetramer staining following antigen stimulation.^43^

The model retained strong performance on all cohorts supporting its robustness across platforms, donors, and species. Detailed cohort descriptions are provided in Supplementary Table 3.

#### (5) Single-cell paired-chain validation

A public paired single-cell dataset containing scRNA-seq and scTCR-seq profiles from 1.1 million T cells was used for paired-chain validation^44^. Of these, 483,838 cells had paired transcriptome and TCR information. The source study annotated 35 T cell populations, including MAIT cells (n = 48,158) and CD8.c16.MAIT.SLC4A10 cells (n = 2,990); iNKT cells were not annotated separately. UcTCRp MAIT and iNKT scores were computed for 431,515 cells with usable TCRβ sequences. Canonical MAIT α-chain-positive cells were defined as those expressing *TRAV1-2* paired with *TRAJ33, TRAJ20* or *TRAJ12*. Three MAIT annotation layers were compared: source transcriptome-based annotation, canonical α-chain annotation and UcTCRp β-chain annotation. For cells not annotated as MAIT in the source dataset, discordant annotation groups were defined as β-chain-only, α+β-supported and α-chain-only. These groups were compared across source cell-type labels and for expression of MAIT-associated genes (*ZBTB16*, *KLRB1* and *SLC4A10*). Cells with iNKT-like UcTCRp scores were additionally assessed for canonical iNKT α-chain usage, defined as *TRAV10* paired with *TRAJ18*.

### Threshold Calibration and Abundance Estimation

To enable quantitative estimation of unconventional T cell subset frequencies from bulk TCRβ repertoires, we leveraged the probabilistic outputs of the UcTCR-Predictor classifier to assign cell-type labels to individual sequences. Each TCRβ sequence was associated with a softmax probability vector across the three classes (MAIT, iNKT, conventional). To achieve high-confidence annotation while minimizing false positives, we optimized a decision threshold for classification.

#### (1) Threshold Optimization

Threshold calibration was performed using an experimentally annotated peripheral blood dataset (Validation Set 2) containing 317 samples from 166 healthy human donors with matched scRNA-seq–derived MAIT and iNKT cell frequencies^42^. Samples containing fewer than 1,000 total TCRβ sequences were excluded from downstream analyses. For each threshold value (ranging from 0.5 to 0.9), we computed the predicted MAIT and iNKT cell proportion (based on the fraction of TCRβ sequences exceeding the threshold for the MAIT class) and compared it to the true cellular abundance across samples. Spearman correlation was calculated between predicted and actual MAIT frequencies at each threshold. Sequences that did not exceed the cutoff for any of the two unconventional classes were assigned to the conventional category. To account for inter-class probability distribution tails, no secondary label reassignment was performed.

#### (2) Simulation of Mixed Repertoires

To evaluate classifier performance in estimating known proportions of unconventional T cells within complex repertoires, we generated in silico mixtures of unconventional and conventional TCRβ sequences. Specifically, for each unconventional subset (MAIT and iNKT), 100 sequences were sampled from the validation set and combined with randomly selected 900 conventional CD4⁺ or CD8⁺ TCRβ sequences, resulting in synthetic repertoires of 1,000 sequences. Mixtures were generated across 50 proportions ranging from 0% to 10% unconventional content. For each mixture, the model was applied to estimate the relative abundance of the corresponding subset, and predictions were compared to the known ground truth to assess quantitative accuracy.

#### (3) Downsampling Robustness Assessment

To assess the effect of sequencing depth on abundance estimation, we randomly selected nine representative healthy TCRβ repertoire from the Emerson et al. cohort^45^ and performed stochastic downsampling at increasing depths (100, 500, 1,000, 5,000, 10,000, 50,000, 100,000, 500,000, 1,000,000 clones). Each depth condition was repeated with 10 replicates. Predicted frequencies stabilized when sample sizes exceeded 1,000 unique TCRβs, with minimal variance across replicates and no bias in under- or overestimation. Below this threshold, predictions became noisier and less reliable. Accordingly, we imposed a minimum sample size cutoff of 1,000 productive TCRβs for inclusion in downstream analyses.

#### (4) Abundance Metric Definitions

For each sample, unconventional T cell subset frequencies were computed using frequency-weighted abundance: The sum of read frequencies for sequences within each class, normalized by the total sequencing depth of the sample. Final abundance estimates were merged with sample metadata including age, sex, disease status, tissue source, and sequencing platform to enable integrative analyses.

### Code availability

The UcTCRp model are available as an open-source Python package at: https://github.com/ZhangLabTJU/UcTCR-Predictor.

## Funding

This work was supported by the National Natural Science Foundation of China (Grant No. 32470688, 32000468), and the Natural Science Foundation of Tianjin (Grant No. 24JCZDJC01400).

## Acknowledgments

We thank Dr. Kunrong Mei, Dr. Tao Li for helpful discussions. We are grateful to all the investigators who made their TCR sequencing datasets publicly available, including through immuneACCESS and other platform.

## Author Contributions

J.Z. supervised and designed this project. L.L., S.S. and Y.D. collected the unconventional TCR data and performed sequence analysis. S.S., Q.X. and Y.D. implemented the classification algorithm, generated the simulation data, and performed method validation on both simulated and experimental data. Y.L., K.W., L.C., H.W., H.W., and C.F. assisted with formal analysis and figure generation. L.L., Y.L., and S.S. wrote the original draft of the manuscript. J.Z. and X.B. performed extensive review and editing of the original manuscript. All co-authors contributed to the research progress discussion and manuscript preparation.

## Declaration of Interests

The authors declare that they have no conflict of interest.

## Supplementary Information

Supplementary materials include the following:

**Supplementary Table 1**: Train, test, and validation dataset sizes of the human ucTCR prediction model.

**Supplementary Table 2**: Train, test, and validation dataset sizes of the mouse ucTCR prediction model.

**Supplementary Table 3**: External validation datasets used in this study

**Supplementary Figure 1**: NMI heatmaps for CDR3βs of varying lengths

**Supplementary Figure 2**: Central CDR3β amino acid usage across different T cell subsets.

**Supplementary Figure 3**: Central CDR3β amino acid usage and position-specific enrichment in human unconventional T cells, stratified by V gene.

**Supplementary Figure 4**: Central CDR3β amino acid usage and position-specific enrichment in mouse unconventional T cells, stratified by V gene.

## Notes

### Competing Interest Statement

The authors have declared no competing interest.

## References

1. Nikolich-Zugich, J., Slifka, M.K., and Messaoudi, I. (2004). The many important facets of T-cell repertoire diversity. Nat Rev Immunol 4, 123–132. 10.1038/nri1292.

2. Pai, J.A., and Satpathy, A.T. (2021). High-throughput and single-cell T cell receptor sequencing technologies. Nat Methods 18, 881–892. 10.1038/s41592-021-01201-8.

3. Mahdy, A.K.H., Lokes, E., Schöpfel, V., Kriukova, V., Britanova, O.V., Steiert, T.A., Franke, A., and ElAbd, H. (2024). Bulk T cell repertoire sequencing (TCR-Seq) is a powerful technology for understanding inflammation-mediated diseases. J Autoimmun 149, 103337. 10.1016/j.jaut.2024.103337.

4. Godfrey, D.I., Uldrich, A.P., McCluskey, J., Rossjohn, J., and Moody, D.B. (2015). The burgeoning family of unconventional T cells. Nat. Immunol. 16, 1114–1123. 10.1038/ni.3298.

5. Kjer-Nielsen, L., Patel, O., Corbett, A.J., Le Nours, J., Meehan, B., Liu, L., Bhati, M., Chen, Z., Kostenko, L., Reantragoon, R., et al. (2012). MR1 presents microbial vitamin B metabolites to MAIT cells. Nature 491, 717–723. 10.1038/nature11605.

6. Borg, N.A., Wun, K.S., Kjer-Nielsen, L., Wilce, M.C.J., Pellicci, D.G., Koh, R., Besra, G.S., Bharadwaj, M., Godfrey, D.I., McCluskey, J., et al. (2007). CD1d–lipid-antigen recognition by the semi-invariant NKT T-cell receptor. Nature 448, 44–49. 10.1038/nature05907.

7. Le Bourhis, L., Martin, E., Péguillet, I., Guihot, A., Froux, N., Coré, M., Lévy, E., Dusseaux, M., Meyssonnier, V., Premel, V., et al. (2010). Antimicrobial activity of mucosal-associated invariant T cells. Nat Immunol 11, 701–708. 10.1038/ni.1890.

8. Lin, Q., Kuypers, M., Philpott, D.J., and Mallevaey, T. (2020). The dialogue between unconventional T cells and the microbiota. Mucosal Immunol 13, 867–876. 10.1038/s41385-020-0326-2.

9. Crosby, C.M., and Kronenberg, M. (2018). Tissue-specific functions of invariant natural killer T cells. Nat Rev Immunol 18, 559–574. 10.1038/s41577-018-0034-2.

10. Dadi, S., Chhangawala, S., Whitlock, B.M., Franklin, R.A., Luo, C.T., Oh, S.A., Toure, A., Pritykin, Y., Huse, M., Leslie, C.S., et al. (2016). Cancer Immunosurveillance by Tissue-Resident Innate Lymphoid Cells and Innate-like T Cells. Cell 164, 365–377. 10.1016/j.cell.2016.01.002.

11. Fan, Q., Nan, H., Li, Z., Li, B., Zhang, F., and Bi, L. (2023). New insights into MAIT cells in autoimmune diseases. Biomedicine & Pharmacotherapy 159, 114250. 10.1016/j.biopha.2023.114250.

12. Van Kaer, L., Parekh, V.V., and Wu, L. (2013). Invariant natural killer T cells as sensors and managers of inflammation. Trends Immunol 34, 50–58. 10.1016/j.it.2012.08.009.

13. Bharadwaj, N.S., and Gumperz, J.E. (2022). Harnessing invariant natural killer T cells to control pathological inflammation. Front. Immunol. 13. 10.3389/fimmu.2022.998378.

14. Godfrey, D.I., Le Nours, J., Andrews, D.M., Uldrich, A.P., and Rossjohn, J. (2018). Unconventional T cell targets for cancer immunotherapy. Immunity 48, 453–473. 10.1016/j.immuni.2018.03.009.

15. Petley, E.V., Koay, H.-F., Henderson, M.A., Sek, K., Todd, K.L., Keam, S.P., Lai, J., House, I.G., Li, J., Zethoven, M., et al. (2021). MAIT cells regulate NK cell-mediated tumor immunity. Nat Commun 12, 4746. 10.1038/s41467-021-25009-4.

16. Laub, A., Rodrigues de Almeida, N., and Huang, S. (2025). Unconventional T cells in anti-cancer immunity. Front. Immunol. 16. 10.3389/fimmu.2025.1618393.

17. Lynch, L., Nowak, M., Varghese, B., Clark, J., Hogan, A.E., Toxavidis, V., Balk, S.P., O’Shea, D., O’Farrelly, C., and Exley, M.A. (2012). Adipose tissue invariant NKT cells protect against diet-induced obesity and metabolic disorder through regulatory cytokine production. Immunity 37, 574–587. 10.1016/j.immuni.2012.06.016.

18. Magalhaes, I., Kiaf, B., and Lehuen, A. (2015). iNKT and MAIT cell alterations in diabetes. Front. Immunol. 6. 10.3389/fimmu.2015.00341.

19. Constantinides, M.G., and Belkaid, Y. (2021). Early-life imprinting of unconventional T cells and tissue homeostasis. Science 374, eabf0095. 10.1126/science.abf0095.

20. Reantragoon, R., Corbett, A.J., Sakala, I.G., Gherardin, N.A., Furness, J.B., Chen, Z., Eckle, S.B.G., Uldrich, A.P., Birkinshaw, R.W., Patel, O., et al. (2013). Antigen-loaded MR1 tetramers define T cell receptor heterogeneity in mucosal-associated invariant T cells. J Exp Med 210, 2305–2320. 10.1084/jem.20130958.

21. Matsuda, J.L., Naidenko, O.V., Gapin, L., Nakayama, T., Taniguchi, M., Wang, C.R., Koezuka, Y., and Kronenberg, M. (2000). Tracking the response of natural killer T cells to a glycolipid antigen using CD1d tetramers. J Exp Med 192, 741–754. 10.1084/jem.192.5.741.

22. Gherardin, N.A., Souter, M.N., Koay, H.-F., Mangas, K.M., Seemann, T., Stinear, T.P., Eckle, S.B., Berzins, S.P., d’Udekem, Y., Konstantinov, I.E., et al. (2018). Human blood MAIT cell subsets defined using MR1 tetramers. Immunol Cell Biol 96, 507–525. 10.1111/imcb.12021.

23. Montoya, C.J., Pollard, D., Martinson, J., Kumari, K., Wasserfall, C., Mulder, C.B., Rugeles, M.T., Atkinson, M.A., Landay, A.L., and Wilson, S.B. (2007). Characterization of human invariant natural killer T subsets in health and disease using a novel invariant natural killer T cell-clonotypic monoclonal antibody, 6B11. Immunology 122, 1–14. 10.1111/j.1365-2567.2007.02647.x.

24. Stubbington, M.J.T., Lönnberg, T., Proserpio, V., Clare, S., Speak, A.O., Dougan, G., and Teichmann, S.A. (2016). T cell fate and clonality inference from single-cell transcriptomes. Nat Methods 13, 329–332. 10.1038/nmeth.3800.

25. Godfrey, D.I., Uldrich, A.P., McCluskey, J., Rossjohn, J., and Moody, D.B. (2015). The burgeoning family of unconventional T cells. Nat Immunol 16, 1114–1123. 10.1038/ni.3298.

26. ElAbd, H., Byron, R., Woodhouse, S., Robinett, B., Sulc, J., Franke, A., Pesesky, M., Zhou, W., Chen-Harris, H., Howie, B., et al. (2024). Seq2MAIT: A Novel Deep Learning Framework for Identifying Mucosal Associated Invariant T (MAIT) Cells. Preprint at Bioinformatics, 10.1101/2024.03.12.584395 https://doi.org/10.1101/2024.03.12.584395.

27. Narayanan, G.A., McLaren, J.E., Meermeier, E.W., Ladell, K., Swarbrick, G.M., Price, D.A., Tran, J.G., Worley, A.H., Vogt, T., Wong, E.B., et al. (2020). The MAIT TCRβ chain contributes to discrimination of microbial ligand. Immunol Cell Biol 98, 770–781. 10.1111/imcb.12370.

28. Dou, Y., Shan, S., and Zhang, J. (2023). UcTCRdb: An unconventional T cell receptor sequence database with online analysis functions. Frontiers in Immunology 14.

29. Grunewald, J., Janson, C.H., and Wigzell, H. (1991). Biased expression of individual T cell receptor V gene segments in CD4+ and CD8+ human peripheral blood T lymphocytes. Eur. J. Immunol. 21, 819–822. 10.1002/eji.1830210342.

30. Klarenbeek, P.L., Doorenspleet, M.E., Esveldt, R.E.E., Schaik, B.D.C. van, Lardy, N., Kampen, A.H.C. van, Tak, P.P., Plenge, R.M., Baas, F., Bakker, P.I.W. de, et al. (2015). Somatic variation of T-cell receptor genes strongly associate with HLA class restriction. PLOS ONE 10, e0140815. 10.1371/journal.pone.0140815.

31. Le Nours, J., Praveena, T., Pellicci, D.G., Gherardin, N.A., Ross, F.J., Lim, R.T., Besra, G.S., Keshipeddy, S., Richardson, S.K., Howell, A.R., et al. (2016). Atypical natural killer T-cell receptor recognition of CD1d–lipid antigens. Nat Commun 7, 10570. 10.1038/ncomms10570.

32. Lepore, M., Kalinichenko, A., Colone, A., Paleja, B., Singhal, A., Tschumi, A., Lee, B., Poidinger, M., Zolezzi, F., Quagliata, L., et al. (2014). Parallel T-cell cloning and deep sequencing of human MAIT cells reveal stable oligoclonal TCRβ repertoire. Nat Commun 5, 3866. 10.1038/ncomms4866.

33. Patel, O., Kjer-Nielsen, L., Le Nours, J., Eckle, S.B.G., Birkinshaw, R., Beddoe, T., Corbett, A.J., Liu, L., Miles, J.J., Meehan, B., et al. (2013). Recognition of vitamin B metabolites by mucosal-associated invariant T cells. Nat. Commun. 4, 2142. 10.1038/ncomms3142.

34. Eckle, S.B.G., Birkinshaw, R.W., Kostenko, L., Corbett, A.J., McWilliam, H.E.G., Reantragoon, R., Chen, Z., Gherardin, N.A., Beddoe, T., Liu, L., et al. (2014). A molecular basis underpinning the T cell receptor heterogeneity of mucosal-associated invariant T cells. J Exp Med 211, 1585–1600. 10.1084/jem.20140484.

35. Li, Y., Girardi, E., Wang, J., Yu, E.D., Painter, G.F., Kronenberg, M., and Zajonc, D.M. (2010). The Vα14 invariant natural killer T cell TCR forces microbial glycolipids and CD1d into a conserved binding mode. J. Exp. Med. 207, 2383–2393. 10.1084/jem.20101335.

36. Lin, Z., Akin, H., Rao, R., Hie, B., Zhu, Z., Lu, W., Smetanin, N., Verkuil, R., Kabeli, O., Shmueli, Y., et al. (2023). Evolutionary-scale prediction of atomic-level protein structure with a language model. Science 379, 1123–1130. 10.1126/science.ade2574.

37. Wu, K.E., Yost, K., Daniel, B., Belk, J., Xia, Y., Egawa, T., Satpathy, A., Chang, H., and Zou, J. (2024). TCR-BERT: learning the grammar of T-cell receptors for flexible antigen-binding analyses. In Proceedings of the 18th Machine Learning in Computational Biology meeting (PMLR), pp. 194–229.

38. Devlin, J., Chang, M.-W., Lee, K., and Toutanova, K. (2019). BERT: Pre-training of Deep Bidirectional Transformers for Language Understanding. Preprint at arXiv, 10.48550/arXiv.1810.04805 https://doi.org/10.48550/arXiv.1810.04805.

39. Glanville, J., Huang, H., Nau, A., Hatton, O., Wagar, L.E., Rubelt, F., Ji, X., Han, A., Krams, S.M., Pettus, C., et al. (2017). Identifying specificity groups in the T cell receptor repertoire. Nature 547, 94–98. 10.1038/nature22976.

40. DeWitt, W.S., III, Smith, A., Schoch, G., Hansen, J.A., Matsen, F.A., IV, and Bradley, P. (2018). Human T cell receptor occurrence patterns encode immune history, genetic background, and receptor specificity. eLife 7, e38358. 10.7554/eLife.38358.

41. Garner, L.C., Amini, A., FitzPatrick, M.E.B., Lett, M.J., Hess, G.F., Filipowicz Sinnreich, M., Provine, N.M., and Klenerman, P. (2023). Single-cell analysis of human MAIT cell transcriptional, functional and clonal diversity. Nat Immunol 24, 1565–1578. 10.1038/s41590-023-01575-1.

42. Terekhova, M., Swain, A., Bohacova, P., Aladyeva, E., Arthur, L., Laha, A., Mogilenko, D.A., Burdess, S., Sukhov, V., Kleverov, D., et al. (2023). Single-cell atlas of healthy human blood unveils age-related loss of NKG2C+GZMB-CD8+ memory T cells and accumulation of type 2 memory T cells. Immunity 56, 2836–2854.e9. 10.1016/j.immuni.2023.10.013.

43. Karnaukhov, V.K., Le Gac, A.-L., Bilonda Mutala, L., Darbois, A., Perrin, L., Legoux, F., Walczak, A.M., Mora, T., and Lantz, O. (2024). Innate-like T cell subset commitment in the murine thymus is independent of TCR characteristics and occurs during proliferation. Proc. Natl. Acad. Sci. U.S.A. 121, e2311348121. 10.1073/pnas.2311348121.

44. Xue, Z., Wu, L., Tian, R., Gao, B., Zhao, Y., He, B., Sun, D., Zhao, B., Li, Y., Zhu, K., et al. (2025). Integrative mapping of human CD8+ T cells in inflammation and cancer. Nat Methods 22, 435–445. 10.1038/s41592-024-02530-0.

45. Emerson, R.O., DeWitt, W.S., Vignali, M., Gravley, J., Hu, J.K., Osborne, E.J., Desmarais, C., Klinger, M., Carlson, C.S., Hansen, J.A., et al. (2017). Immunosequencing identifies signatures of cytomegalovirus exposure history and HLA-mediated effects on the T cell repertoire. Nat Genet 49, 659–665. 10.1038/ng.3822.

46. Chandra, S., Ascui, G., Riffelmacher, T., Chawla, A., Ramírez-Suástegui, C., Castelan, V.C., Seumois, G., Simon, H., Murray, M.P., Seo, G.-Y., et al. (2023). Transcriptomes and metabolism define mouse and human MAIT cell populations. Sci. Immunol. 8, eabn8531. 10.1126/sciimmunol.abn8531.

47. Gherardin, N.A., Keller, A.N., Woolley, R.E., Le Nours, J., Ritchie, D.S., Neeson, P.J., Birkinshaw, R.W., Eckle, S.B.G., Waddington, J.N., Liu, L., et al. (2016). Diversity of T Cells Restricted by the MHC Class I-Related Molecule MR1 Facilitates Differential Antigen Recognition. Immunity 44, 32–45. 10.1016/j.immuni.2015.12.005.

48. Awad, W., Meermeier, E.W., Sandoval-Romero, M.L., Le Nours, J., Worley, A.H., Null, M.D., Liu, L., McCluskey, J., Fairlie, D.P., Lewinsohn, D.M., et al. (2020). Atypical TRAV1-2− T cell receptor recognition of the antigen-presenting molecule MR1. J Biol Chem 295, 14445–14457. 10.1074/jbc.RA120.015292.

49. Mhanna, V., Bashour, H., Lê Quý, K., Barennes, P., Rawat, P., Greiff, V., and Mariotti-Ferrandiz, E. (2024). Adaptive immune receptor repertoire analysis. Nat Rev Methods Primers 4, 6. 10.1038/s43586-023-00284-1.

50. Seay, H.R., Yusko, E., Rothweiler, S.J., Zhang, L., Posgai, A.L., Campbell-Thompson, M., Vignali, M., Emerson, R.O., Kaddis, J.S., Ko, D., et al. (2016). Tissue distribution and clonal diversity of the T and B cell repertoire in type 1 diabetes. JCI Insight 1. 10.1172/jci.insight.88242.

