## Supplementary Materials for "UcTCRp: a TCRβ-based framework for quantitative MAIT- and iNKT-associated repertoire-state profiling"

Jian Zhang *et al.*

### The PDF file includes:

- Supplementary Figures 1 to 4
- Supplementary Tables 1 to 3
- References

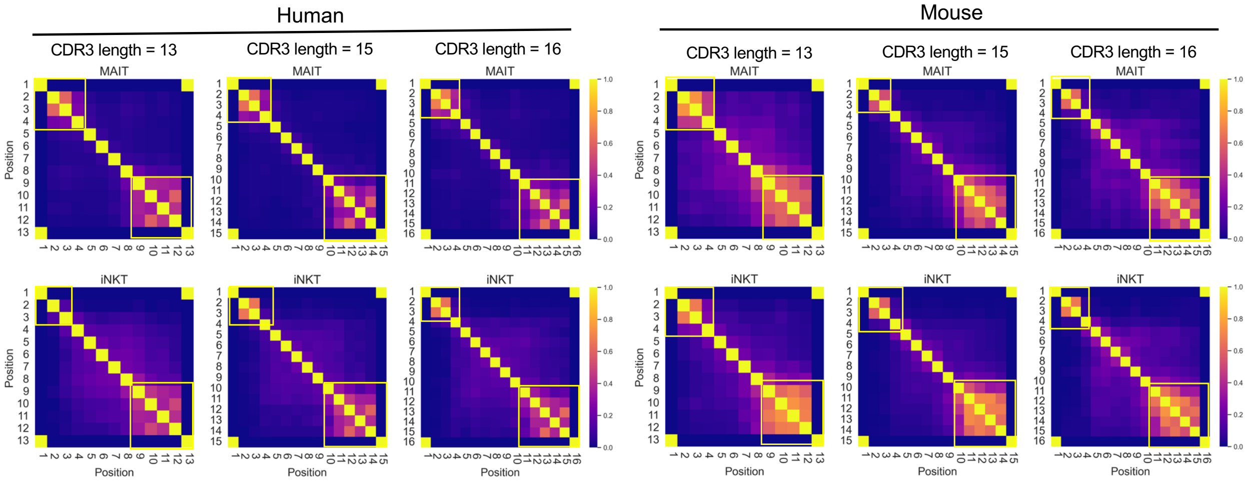


**Supplementary Figure 1. NMI heatmaps for CDR3βs of varying lengths.** Position-wise normalized mutual information (NMI) across CDR3β sequences (lengths 13, 15, 16) in human and mouse MAIT and iNKT cells, illustrating germline-constrained terminal regions versus recombination-driven central diversity.

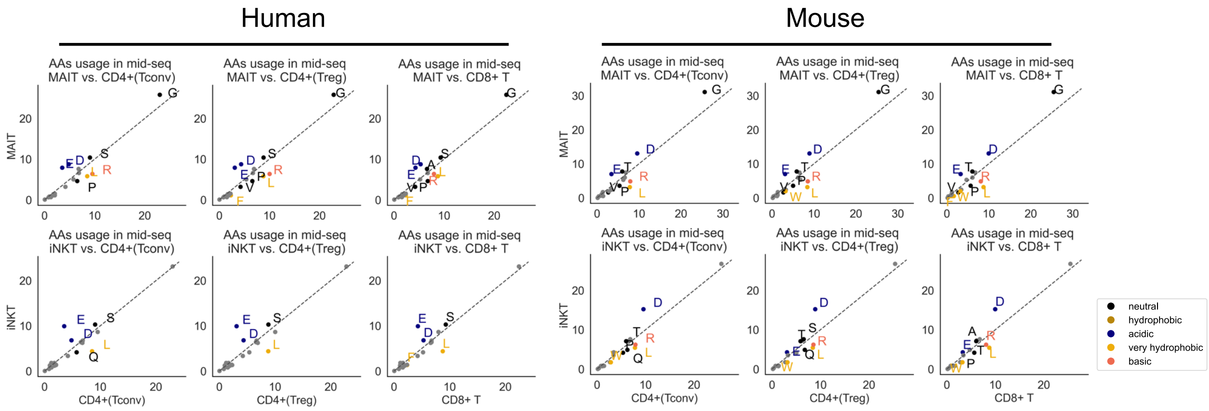


**Supplementary Figure 2. Central CDR3β amino acid usage across different T cell subsets.** Amino acid usage profiles in the central "mid-seq" region of CDR3β sequences (positions excluding the first 4 and last 6 residues) across unconventional (MAIT, iNKT) and conventional ($CD4^+$ Tconv, $CD4^+$ Treg, $CD8^+$) T cell subsets.

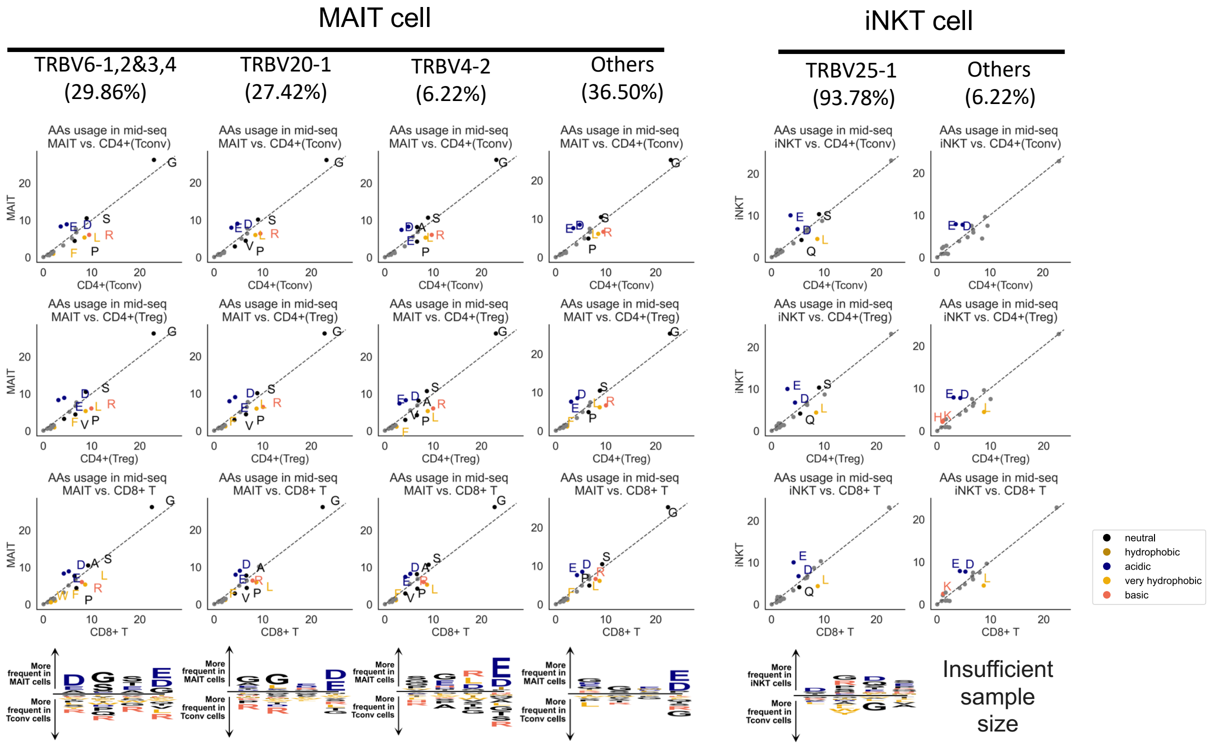


**Supplementary Figure 3. Central CDR3β amino acid usage and position-specific enrichment in human unconventional T cells, stratified by V gene.** Amino acid usage profiles in the central "mid-seq" region of CDR3β sequences (excluding the first 4 and last 6 residues) across unconventional (MAIT, iNKT) and conventional (CD4$^{+}$ Tconv, CD4$^{+}$ Treg, CD8$^{+}$) T cell subsets in human. The bottom panels show position-specific enrichment of amino acid residues at positions 5–8 in fixed-length (14-aa) CDR3β sequences. Analyses are stratified by V gene usage to control for germline-encoded influences, revealing conserved biochemical biases that persist independent of V gene identity.

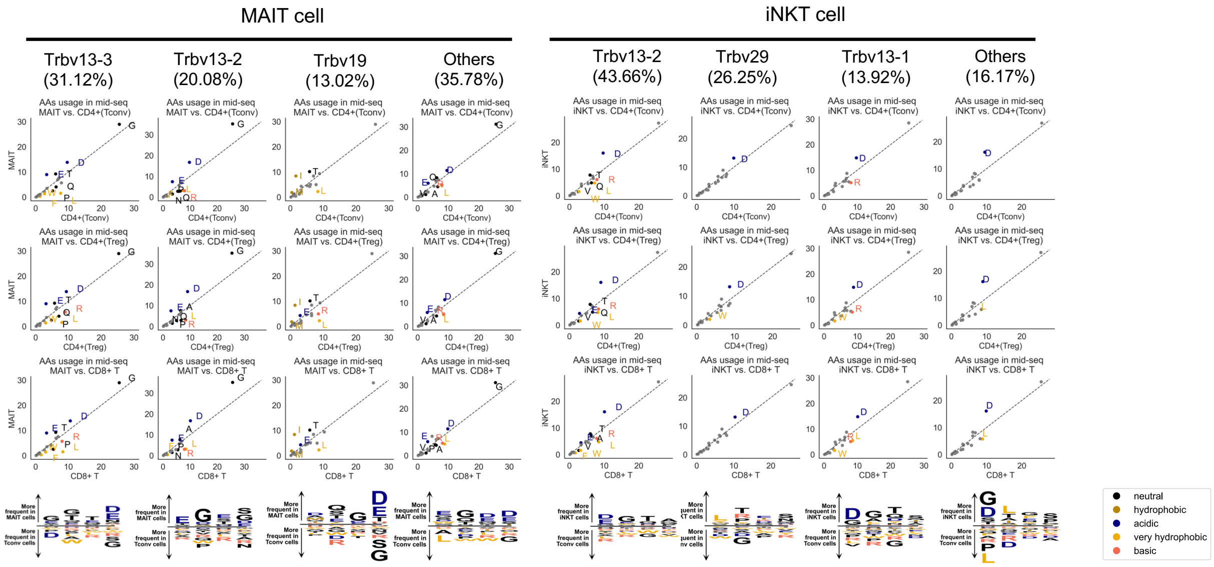


**Supplementary Figure 4. Central CDR3β amino acid usage and position-specific enrichment in mouse unconventional T cells, stratified by V gene.** Amino acid usage profiles in the central "mid-seq" region of CDR3β sequences (excluding the first 4 and last 6 residues) across unconventional (MAIT, iNKT) and conventional (CD4+ Tconv, CD4+ Treg, CD8+) T cell subsets in mice. The bottom panels show position-specific enrichment of amino acid residues at positions 5–8 in fixed-length (14-aa) CDR3β sequences. Analyses are stratified by V gene usage to control for germline-encoded influences, revealing conserved biochemical biases that persist independent of V gene identity.

**Supplementary Table 1. Train, test, and validation dataset sizes of the HUMAN ucTCR prediction model.**

| Dataset | MAIT | iNKT | Conventional T (CD4+, CD8+) | Total |
|---|---|---|---|---|
| **Train** | 18,014 | 1950 | 198,860 | 218,824 |
| **Test** | 2003 | 219 | 22,092 | 24,314 |
| **Validation** | 100 | 100 | 100 | 300 |
| **Total** | 20,117 | 2,269 | 221,052 | 243,438 |

**Supplementary Table 2. Train, test, and validation dataset sizes of the MOUSE ucTCR prediction model.**

| Dataset | MAIT | iNKT | Conventional T (CD4+, CD8+) | Total |
|---|---|---|---|---|
| **Train** | 1,492 | 1,988 | 34,790 | 38,270 |
| **Test** | 194 | 222 | 3,837 | 4,253 |
| **Validation** | 100 | 100 | 100 | 300 |
| **Total** | 1,786 | 2,310 | 38,727 | 42,823 |

**Supplementary Table 3. External validation datasets used in this study**

| Dataset | Source | Species | Tissue | Cell Type | No. of TCRβs | Subset identification method | Accession |
|---|---|---|---|---|---|---|---|
| Validation Set 1 | Garner et al., 2023[1] | Human | Liver, blood | MAIT | 17,772 | Antigen simulation | GEO: GSE194189 |
| Validation Set 2,3 | Terekhova et al, 2023[2] | Human | Blood | MAIT, iNKT | 17,340/324 | Single-cell RNA-seq | Synapse: syn49637038 |
| Validation Set 4 | Karnaukh ov et al, 2024[3] | Mouse | Thymu s | MAIT | 845 | Antigen simulation | GEO: GSE236666 |

### REFERENCES


1. Garner, L.C., Amini, A., FitzPatrick, M.E.B., Lett, M.J., Hess, G.F., Filipowicz Sinnreich, M., Provine, N.M., and Klenerman, P. (2023). Single-cell analysis of human MAIT cell transcriptional, functional and clonal diversity. Nat Immunol *24*, 1565–1578. https://doi.org/10.1038/s41590-023-01575-1.

2. Terekhova, M., Swain, A., Bohacova, P., Aladyeva, E., Arthur, L., Laha, A., Mogilenko, D.A., Burdess, S., Sukhov, V., Kleverov, D., et al. (2023). Single-cell atlas of healthy human blood unveils age-related loss of NKG2C+GZMB−CD8+ memory T cells and accumulation of type 2 memory T cells. Immunity *56*, 2836-2854.e9. https://doi.org/10.1016/j.immuni.2023.10.013.

3. Karnaukhov, V.K., Le Gac, A.-L., Bilonda Mutala, L., Darbois, A., Perrin, L., Legoux, F., Walczak, A.M., Mora, T., and Lantz, O. (2024). Innate-like T cell subset commitment in the murine thymus is independent of TCR characteristics and occurs during proliferation. Proc. Natl. Acad. Sci. U.S.A. *121*, e2311348121. https://doi.org/10.1073/pnas.2311348121.
